# Drought resistance and improved yield result from modified malate metabolism in guard and vascular companion cells

**DOI:** 10.1101/2024.06.24.600218

**Authors:** Pablo Oitaven, María F. Guindón, Gabriela L. Müller, Ezequiel Margarit, Carolina Saper, María Sol Srebot, Ying Fu, Karuna Verma, Vera Wewer, Sabine Metzger, María V. Lara, Gonzalo Martin Estavillo, Veronica G. Maurino, María F. Drincovich

## Abstract

Drought is a major threat to food security. Water loss through stomata is an inevitable consequence of CO_2_ uptake, and water deficit inhibits plant growth, making it challenging to develop drought-resistant strategies without compromising yield. Here, we generated tobacco plants expressing a maize NADP-dependent malate decarboxylating enzyme in stomata and vascular cells (ME plants), which show higher seed yield and faster maturation compared to wild-type (WT) plants under normal irrigation and after drought. While WT plants die after 45 days of drought, ME plants survive without any adverse effects on seed production. In addition, ME plants exhibit improved photosynthetic efficiency despite reduced stomatal conductance and changes in stem morphology, which are likely related to their ability to withstand drought. We propose that enhanced C_4_-like biochemistry in cells surrounding the vascular system and increased sugar export likely compensated for the reduced stomatal conductance in ME plants. The study demonstrates that cell-targeted metabolic modifications can avoid pleiotropic effects and facilitate the stacking of beneficial traits to improve crop design.

**Significance Statement:** Drought is one of the biggest threats to global food security, and its impact on crop yield is expected to worsen due to climate change. Traditionally, drought resistance has often come at the expense of yield, creating a negative trade-off. However, we present here a promising solution to this challenge. We have developed a novel approach that successfully uncouples the negative balance between drought resistance and yield. By introducing a maize enzyme into specific tobacco cells, we have created drought-resistant plants with faster growth and higher seed yield. Most importantly, after prolonged drought, while the wild type dies, the modified plants maintain their high yield. This technology paves the way for greater food security and resilience to climate change.

## Introduction

The expected increase in the frequency and intensity of extreme weather events such as droughts and heat waves over the next century (1) will increasingly affect food security. This creates an urgent need to develop novel strategies to produce high-yielding crops with stable productivity under worsening climatic conditions (2, 3). The rational development of climate-smart and productive crops requires in-depth knowledge of all aspects that underpin crop productivity, including light capture, CO_2_ fixation, carbohydrate production in source organs, and carbon allocation to sink tissues (4–6).

The most damaging perturbation of the photosynthetic process and the long-distance transport to sink tissues is water shortage (7). Accordingly, drought causes the most significant losses in annual crop yield (8). In the source tissues of terrestrial plants, CO_2_ uptake and water loss occur primarily through stomata; therefore, water loss from aerial tissues during CO_2_ uptake is inevitable (9). In response to drought, plants close stomata, resulting in reduced CO_2_ uptake (10), so drought-tolerant plants with reduced stomatal conductance often show reduced carbon uptake (11, 12). Recent approaches that replace steady-state conductance engineering with alterations of stomatal speed and responsiveness have proved more appropriate to overcome the physical constraints of CO_2_ and water exchange (reviewed in (13)).

Drought also inhibits plant growth by repressing cell division and proliferation (14). As a result, many drought-tolerant plants that overexpress stress-inducible genes exhibit growth retardation and lower yields even in well-watered conditions (15, 16). Attempts to overcome the trade-off between drought resistance and growth have included the stacking of drought-tolerance with growth-enhancing genes (15) or cell-localized overexpression approaches (17).

One promising way to improve drought resistance is to alter intracellular concentrations of the organic acid malate. Reducing malate concentrations in guard cells, which surround and control stomatal movements, can reduce stomatal aperture through a direct osmotic effect or by reducing the activation of chloride channels (18). We therefore introduced a maize NADP-dependent malic enzyme (ZmnpNADP-ME, (19)) under the control of the potassium channel 1 (KAT1) promoter (20) into the guard cells of *Nicotiana tabacum* to catalyze the irreversible malate oxidative decarboxylation to pyruvate, CO_2_, and NADPH. Three independent *KAT1::ZmnpNADP-ME* transgenic tobacco lines (ME plants) showed not only reduced stomatal aperture, but also an unexpected increase in sugar export from the leaves compared to the wild type (WT) (21), which may result from ZmnpNADP-ME expression from the KAT1 promoter in vascular companion cells (22).

Here, we show that the ME plants are highly resistant to prolonged drought. In contrast to other drought-tolerant plants produced to date, the resistance of ME plants to water shortage is associated with a higher seed yield than in the WT under normal irrigation, and the full yield is achieved in a shorter time. Strikingly, this high seed yield is maintained even after recovery from prolonged severe drought. Thus, we find that the strategy of targeting maize NADP-ME expression to both guard and vascular companion cells can overcome the trade-off between drought resistance and yield.

## Results

### Plants with increased malate metabolism (ME plants) can recover from prolonged drought

To test the response of the three independent *KAT1::ZmnpNADP-ME* tobacco lines ME1, ME3, and ME4 (ME plants) to drought, at least 10 plants of each genotype were grown, together with the WT, under well-watered conditions (90 % Field Capacity, FC) for 60 days and then watering was stopped for either 30 (30D) or 45 days (45D). The plants were then rewatered to maturity to assess their recovery (Fig. 1). Although both WT and ME plants showed clear signs of drought stress after 30D, the transgenic lines were less affected with milder wilting symptoms than the WT (30D, Fig. 1A). When we rewatered the plants at 30D, the leaves of the transgenic lines showed signs of recovery (less wilting than the WT) after just one day (30D1R, Fig. 1A). After a further seven days of rewatering (30D8R), significantly more leaves (p˂0.001) had recovered in the transgenic lines (almost 45% of all leaves) than in the WT (less than 32% of all leaves) (30D8R, Fig. 1A).

**Fig. 1.**
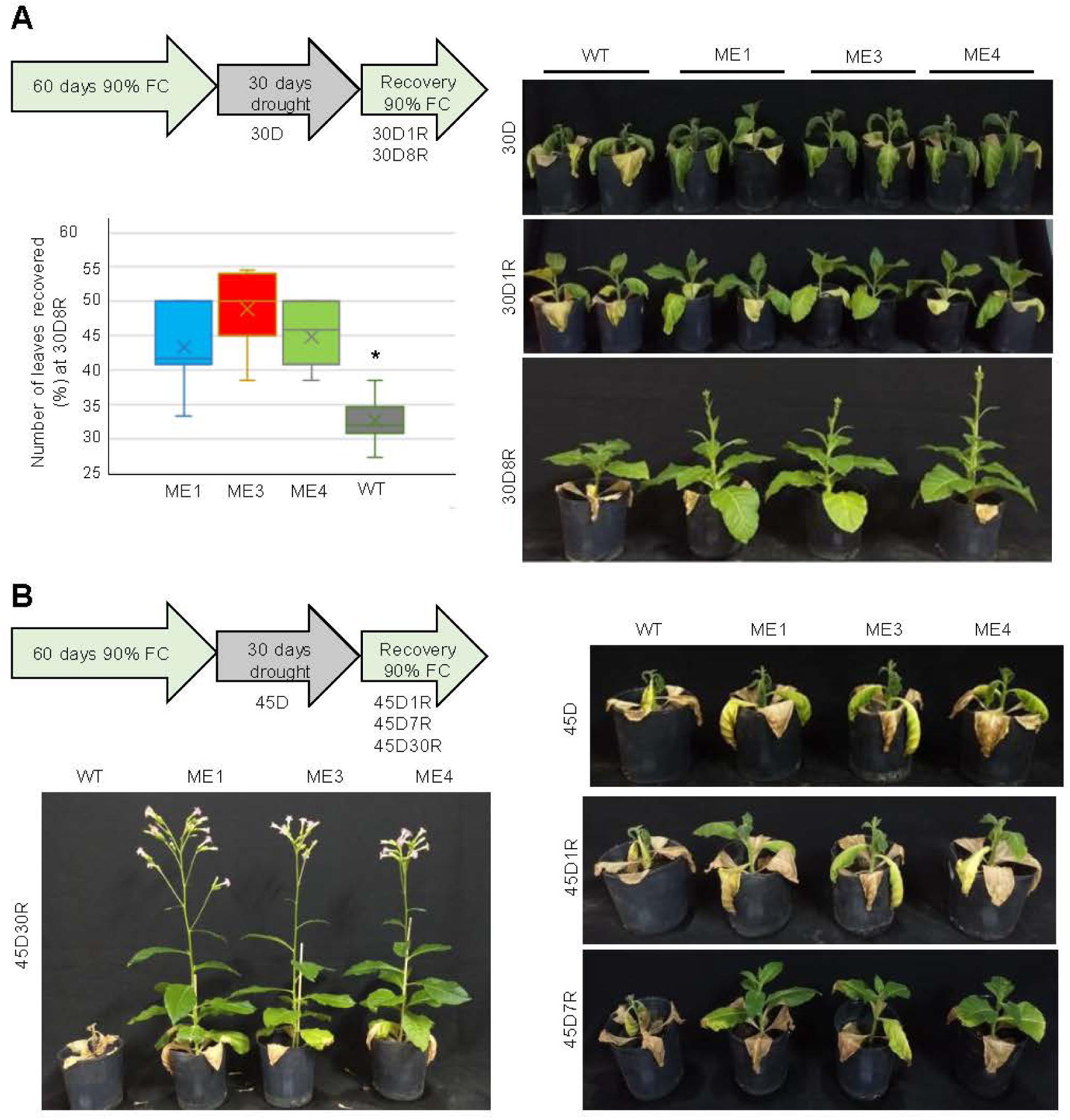
Drought and recovery treatments. Tobacco WT and the transgenic lines ME1, ME3, and ME4 were grown at 90% FC for 60 days. Watering was stopped for 30 (**A**) or 45 days (**B**), followed by rewatering at 90% FC. A. Images of plants exposed to drought for 30 days (30D) and after rewatering for 1 (30D1R) and 8 (30D8R) days. The percentage of leaves recovered at 30D8R out of the total number of leaves of each plant is shown on the left. Measurements were conducted in 10 plants per genotype. * indicates statistically significant difference (*p* value <0.001) in Kruskal-Wallis tests. Four independent biological replicates were conducted from 2018 to 2021 with similar results. Images are from 2018. B. Images of plants exposed to drought for 45 days (45D) and after rewatering for 1 (45D1R), 7 (45D7R), and 30 days (45D30R). Three independent biological replicates were conducted from 2018 to 2021 with similar results.

After 45 days of drought, both WT and ME lines showed severe wilting (Fig. 1B). After only one day of rewatering, the ME plants showed signs of recovery (45D1R, Fig. 1B) and visibly more turgid leaves after eight days (45D8R, Fig. 1B). Most importantly, after 30 days of rewatering (45D30R), 88% of the transgenic lines (8 out of 9 plants in two different biological repetitions) recovered and flowered, while no WT plant survived the treatment (Fig. 1B).

### ME lines flower earlier and have higher seed yield when well-watered and after drought

The ME plants flower and produce mature seeds earlier than the WT when grown at 90% FC and after 30D followed by rewatering (30D+R) (Table S1). On average, the three ME lines flowered and produced mature seeds about 20 days earlier than the WT after recovery from 30D, as observed for plants grown at 90% FC (Table S1).

We measured the aerial vegetative (leaves and stem; Table S2) and reproductive (seeds; Fig. 2 and Table S3) biomass of the WT and ME lines at the end of the life cycle at 90% FC and after recovery from 30 (30D+R) and 45 (45D+R) days of drought. Considering the accelerated development program of the ME lines (Table S1), the biomass values of the ME plants were achieved in a shorter time (0.79 to 0.87 times) than those of the WT plants. The WT plants produced approximately 1.5 times more vegetative aerial dry biomass (dry weight, DW) than the ME plants both under well-watered conditions and at 30D+R (Table S2). However, while no viable WT plant recovered at 45D+R (Fig. 1B), the corresponding ME plants had similar vegetative biomass as at 30D+R (Table S2).

**Fig. 2.**
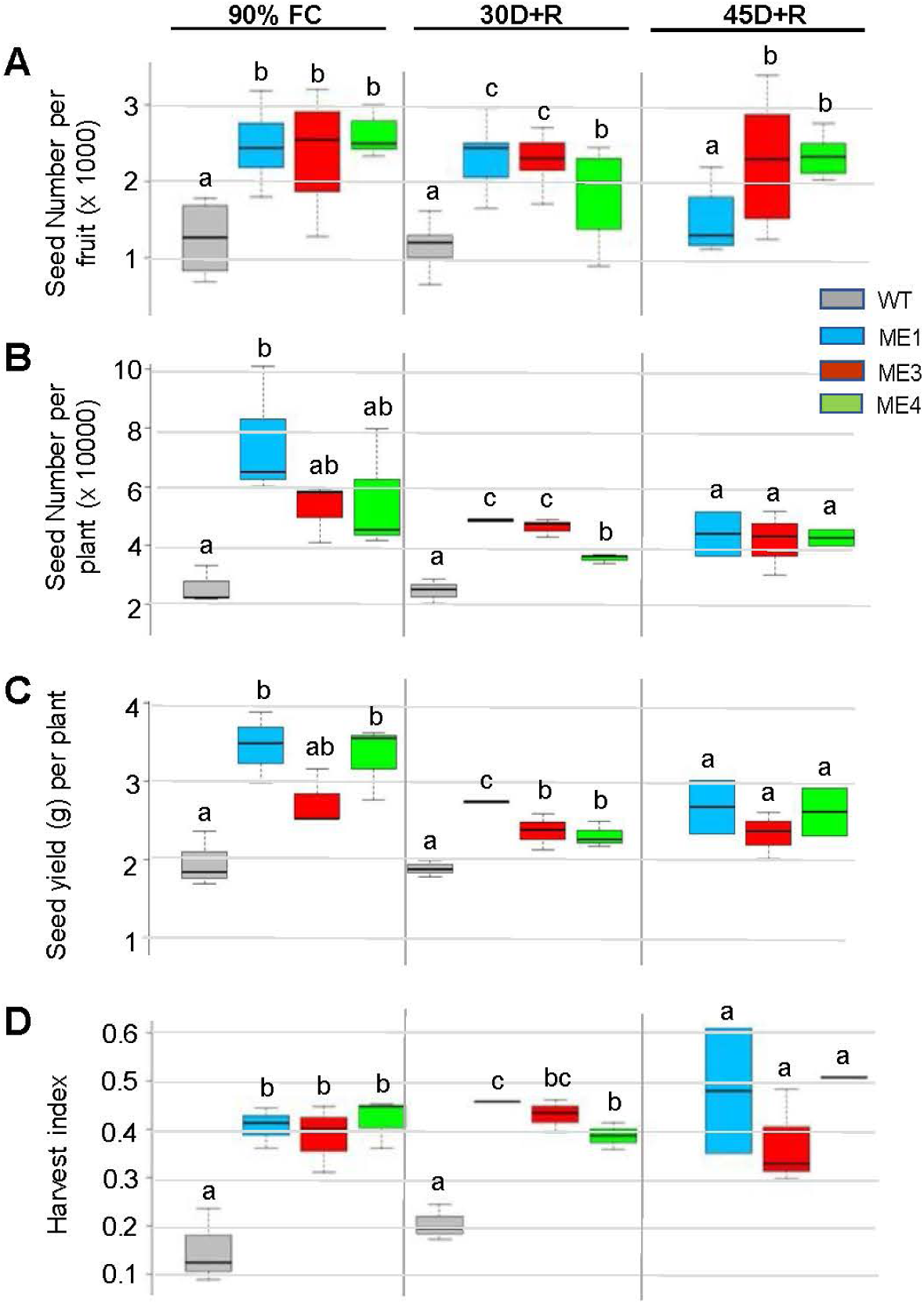
Seed yield and harvest index under control conditions (90 % FC) and after recovery from 30 (30D+R) and 45 days of drought (45D+R). Total reproductive biomass was measured in 2 independent biological replicates with similar results, using at least 3 plants of each genotype. The results are from the experiment in 2021. **A.** Number of seeds per fruit. **B.** Number of seeds per plant. **C.** Seed yield (in g) per plant. **D.** Harvest index. Different letters indicate significant differences (*p*<0.05) according to Tukeýs test of ANOVA or the Kruskal-Wallis test (number of seeds per fruit at 30D+R, and harvest index at 90% FC and 30D+R). The WT did not recover after 45D+R (Fig. 1B). Table S3 shows the full reproductive biomass data.

In terms of reproductive biomass, the number of fruits per plant was not significantly different between the ME and WT plants, either at 90% FC or at 30D+R (Table S3). However, the number of seeds per fruit was 1.7-2.1 times higher in the ME plants than in the WT, both under control conditions and at 30D+R (Fig. 2A), resulting in 1.7-2.9 times more seeds per plant than in the WT (Fig. 2B). Most importantly, while the WT did not survive after 45D, the corresponding ME plants showed almost the same number of fruits per plant and seeds per fruit and plant as after recovery from 30D (Fig. 2A, 2B, and Table S3).

The seed weight of the ME plants was between 1.1 and 1.5 times lower than that of the WT, either under control conditions or at 30D+R (Table S3). However, due to the higher number of seeds, the seed yield (g of seeds per plant) of the ME lines increased by 1.4 to 1.8 times compared to the WT, either under 90% FC or at 30D+R (Fig. 2C). In addition, the seed yield of the ME lines at 45D+R was similar to their seed yield at 30D+R (Fig. 2C). The harvest index (HI), defined as the ratio of seed yield to vegetative plant biomass (Table S2), was 2.3 to 2.8 times higher in the ME plants than in the WT, either under control conditions or at 30D+R (Fig. 2D). Furthermore, the ME lines maintained their high HI at 45D+R (Fig. 2D).

As the seed weight of the ME plants was lower than that of the WT (Table S3), we analyzed their composition and size (Table S4). The water, starch, triacylglycerol, and protein contents between the ME lines and the WT were not statistically significantly different under control conditions or at 30D+R (Table S4). However, the seeds of the ME lines were slightly smaller than those of the WT (Table S4). After recovery from 30D or 45D, seeds were comparable in composition and size to those of plants grown under control conditions, except for the WT at 45D, for which no plant reached maturity (Table S4).

### Stems of ME plants differ in morphology from WT

Stems connect the plant organs and are responsible for the long-distance transport of water, photosynthates, and many other compounds. During severe drought, the stems are prone to cavitation due to bubbles trapped within xylem conduits, preventing or delaying recovery when irrigation is restored (23). Given the rapid recovery of the ME lines after rewatering following drought (Fig. 1), we further analyzed the stem morphology of the ME plants.

Stem light microscopy sections revealed differences in the distribution of the stem tissues (Fig. 3). For 60-day-old plants at 90% FC, the pith cross-sectional length was between 1.2 and 1.4 times greater in the ME lines than in the WT, resulting in 1.2, 1.9, and 1.7-times greater pith areas in the stems of ME1, ME3, and ME4, respectively, than in the WT (Table S5). In addition, the ME lines had a smaller (0.3 to 0.7 times) width of xylem tissue than the WT, resulting in a smaller (0.4 to 0.6 times) area occupied by the xylem compared to the WT (Table S5). Overall, at 90% FC, the pith occupied approximately 13% of the WT stem area, whereas the xylem occupied nearly 15%. In the ME lines, between 20 and 31% of the stem area was occupied by the pith, while the xylem occupied 8-10% (Fig. 3A). Furthermore, the reduction in the xylem area in the ME lines was accompanied by an increase in the number of small-diameter xylem vessels and a reduction in the number of the large ones compared to the WT (Fig. 3A).

**Fig. 3.**
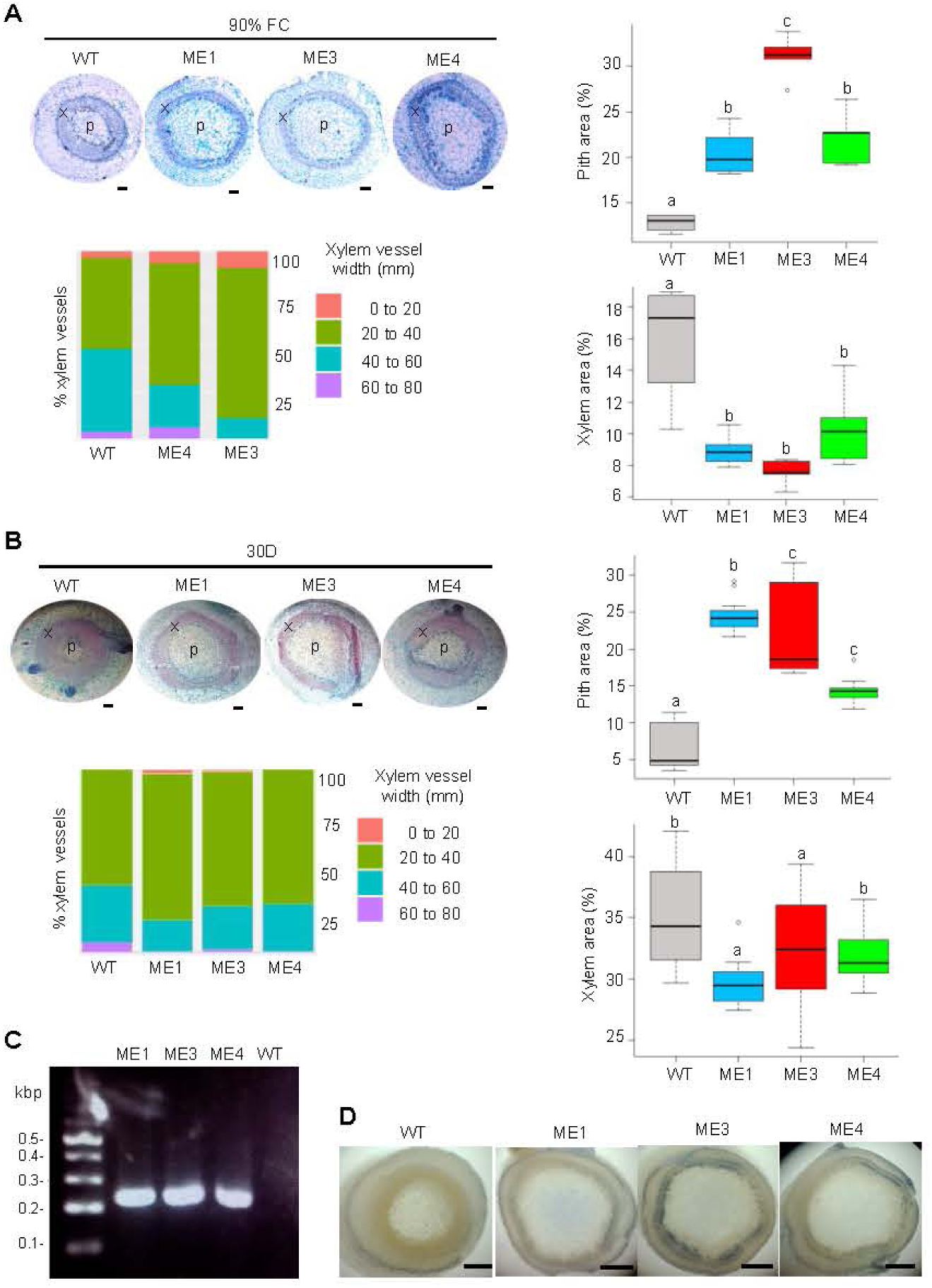
Morphology and *in situ* NADP-ME activity of stems from WT and ME plants. Representative light microscopic stem cross sections, % of xylem vessels as a function of diameter (0-20, 20-40, 40-60, and 60-80 µm), and % of pith and xylem area of stems from plants grown at 90% FC for 60 days (**A**) and after 30 days of drought (30D) (**B**). **C.** Stem cross-sections were used to calculate pith, xylem, and cortex diameters and areas (Table S5). Different letters indicate significant differences (*p*<0.05) according to Tukeýs test of ANOVA or the Kruskal-Wallis test (xylem area at 30D). p: pith, x: xylem. **D.** Detection of the *ZmnpNADP-ME* transcript (230 bp) in stems by RT-PCR. **E.** *In situ* NADP-ME activity staining in stem cross sections of 60-day-old plants at 90% FC. The bars correspond to 1 mm.

At 30D, the pith main cross-sectional length pith was 1.5 to 2 times greater in the ME lines than in the WT, resulting in 2.5 to 3.8 times greater areas occupied by the pith (Fig. 3B, Table S5). In addition, the ME lines had a significantly smaller width of the xylem tissue and a smaller xylem area than the WT (Fig. 3B, Table S5). Overall, at 30D, approximately 6% of the WT stem area was occupied by pith, whereas approximately 35 % was occupied by xylem. Conversely, 14 to 24% of the stem area was occupied by pith, and 30 to 32% by xylem in the ME lines (Fig. 3B). As also observed at 90% FC (Fig. 3A), the ME lines at 30D showed an increase in the number of small-diameter xylem vessels and a decrease in the number of large ones compared to the WT (Fig. 3B).

Given the significant morphological differences between the WT and ME plants, we examined the expression of *ZmnpNADP-ME* and the enzymatic activity in the stems. Apart from the surprising detection of *ZmnpNADP-ME* transcripts in the stems of the ME lines (Fig. 3C), NADP-ME activity was found near the vasculature of the ME lines grown at 90% FC, whereas it was negligible in the stems of WT plants (Fig. 3D).

### ME lines have reduced stomatal conductance independent of ABA

To investigate the potential stress response mechanism in the ME lines, we analyzed stomatal conductance dynamics at 90% FC, during drought stress (30D), and after rewatering for three (30D3R) and seven (30D7R) days. The stomatal conductance (gs) of the ME lines were almost 2-fold lower than those of the WT in developed leaves at 90% FC (about 200 mmol m^−2^ s^−1^ for the WT versus an average of 100 mmol m^−2^ s^−1^ for the transgenic lines; Fig. 4A). After 30D, the gs of WT and ME plants was drastically reduced and reached similar values in all genotypes (about 48 mmol m^−2^ s^−1^; Fig. 4A). After three days of rewatering (30D3R), the gs increased up to similar levels (about 100 mmol m^−2^ s^−1^) in WT and ME lines (Figu. 4A). After seven days of rewatering (30D7R), the gs of the WT was slightly higher than that of the ME lines and similar to the pretreatment levels (Fig. 4A).

**Fig. 4.**
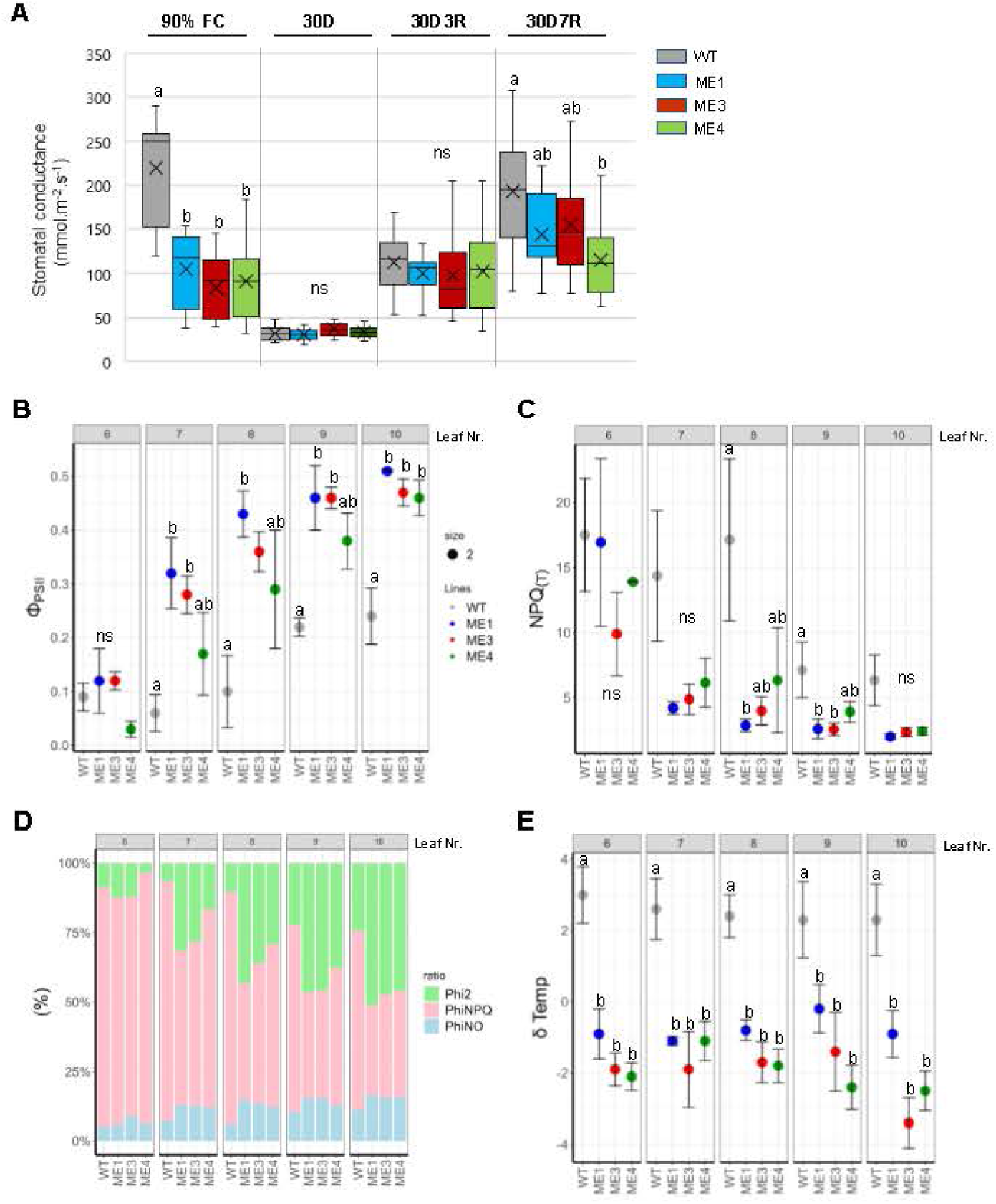
Stomatal conductance and chlorophyll fluorescence measurements. **A.** Stomatal conductance of the fifth or sixth leaf after 4-6 h of illumination under control conditions (90% FC; 60-day-old plants), after 30 days of drought (30D), and after 3 (30D3R) and 7 (30D7R) days of rewatering. Different letters indicate significant differences at the level of 0.001 according to Tukeýs test of ANOVA (90% FC) or at the level of 0.002 according to Kruskal-Wallis test (30D7R). ns: not significant. **B to E:** Chlorophyll fluorescence measurements at 30D of individual leaves 6 to 10 of three different pants of each genotype using the MultispeQ v2.0 sensor (PhotosynQ platform Project ID 7925). The leaf number (6 to 10) is indicated at the top of each plot. **B.** Quantum yield of photosystem PSII (Φ_PSII_). **C.** Theoretical non-photochemical quenching (NPQ_(T)_). **D.** Energy distribution (%) in photochemical (Φ_PSII_, green), non-photochemical (Φ_NPQ_, pink), and non-regulatory quenching (Φ_NO_, light blue). **E.** Difference between leaf and ambient temperature (δ Temp). Different letters indicate significant differences in pairwise comparisons at a 0.05 significance level according to Tukeýs test of ANOVA. ns: not significant.

Abscisic acid (ABA), specifically the isomer (+)-*cis, trans*-ABA (24), mediates the decrease in stomatal conductance (25). We measured ABA levels at 90% FC and after 7 days of drought in leaves of the WT and ME lines (Table S6). At 90% FC, (+)-*cis, trans*-ABA levels were similar between the WT and ME lines although the gs of the ME lines was lower than that of the WT (Table S6). After 7 days of drought (7D), (+)-*cis, trans*-ABA levels increased 30-fold in WT and 12- to 16-fold in ME plants, accompanied by a drastic decrease in gs in both WT and ME plants (Table S6). Levels of (+)-*trans, trans*-ABA were approximately 10-fold lower than those of (+)-*cis, trans*-ABA at 90% FC and increased only 3-fold after 7D in all genotypes, confirming that it is an isomer that is generally present at low levels in plant tissues (26).

### ME lines maintain high photosynthesis during drought

We performed chlorophyll fluorescence measurements on the youngest fully developed leaf (number 5) of 60-day-old WT and ME lines at 90% FC before watering was stopped. Despite the lower gs of the ME lines compared to WT (Fig. 4A), there were no significant differences between WT and ME plants in the photosystem PSII quantum yield (Φ_PSII_), non-regulated energy dissipation (Φ_NO_), non-photochemical quenching (Φ_NPQ_), linear electron flow (LEF), relative chlorophyll content (SPAD), and the difference between leaf and ambient temperature (δ Temp) (Table S7).

At 30D, we performed chlorophyll fluorescence measurements on leaves 6 to 10, counting from the bottom (Dataset S1). In the WT, younger leaves 10 and 9 showed higher Φ_PSII_ (Fig. 4B) and LEF (Fig. S1A) than older leaves 8 to 6. In the ME lines, Φ_PSII_ and LEF were higher in leaves 7 to 10 than in leaf 6. In addition, at 30D, leaves 7 to 10 of the ME lines had significantly higher Φ_PSII_ and LEF than the corresponding WT leaf. The Φ_PSII_ and LEF values in leaf 10 of the ME lines reached maximum values, with an almost 2-fold increase compared to the WT (Fig. 4B, Fig. S1A). The relative chlorophyll content (SPAD) values were not significantly different when comparing the corresponding leaves of WT and ME lines at 30D (Fig. S1B), indicating that the increase in LEF was independent of SPAD.

Conversely, older leaves of 30D plants had higher Φ_NPQ_ (Dataset S1) and theoretical non-photochemical quenching (NPQ_(T)_) than younger leaves (Fig. 4C). In WT, leaves 6 to 8 had higher NPQ_(T)_ than leaves 9 and 10, whereas in ME lines only leaf 6 had higher values than all other younger leaves (Fig. 4C). In addition, WT plants had higher Φ_NPQ_ (Dataset S1) and NPQ_(T)_ values (up to 3-fold) in leaves 8 and 9 than the corresponding leaves of the ME lines (Fig. 4C).

Overall, at 30D, the dissipative photochemical (Φ_PSII_) and non-photochemical (Φ_NPQ_ and Φ_NO_) processes for the energy absorbed by PSII differed between the WT and the ME lines: in the younger leaves (7 to 10), the ME lines dissipated a more significant fraction of the energy absorbed by PSII by photochemical quenching (higher Φ_PSII_ than WT), whereas in WT leaves dissipation by NPQ was predominant (higher Φ_NPQ_ than ME lines), with a low contribution from Φ_NO_ (Fig. 4D). Φ_NO_ generally contributes to a small fraction of non-photochemical quenching compared to Φ_NPQ_ and it was already shown to remain stable after five days of drought in tobacco leaves (27).

We measured leaf temperature and calculated the difference between leaf and ambient temperature (δ Temp). Although there were no significant differences in δ Temp between 60-day-old WT and ME lines at 90% FC (all leaves were cooler than ambient; Table S7), significant differences were observed after 30D (Fig. 4E). In this case, all measured leaves (6 to 10) from the WT plants were about 3° C warmer than ambient temperature, whereas those of the ME lines were significantly cooler, with δ Temp values similar to those observed in all genotypes at 90% FC (Fig. 4E, Table S7).

### Higher levels of photosynthesis-related proteins are present in ME lines than WT during drought

The ME lines differ from the WT in anatomical and physiological aspects, including an enhanced resistance to drought. To identify molecular processes that might be involved in these differences, we performed a comparison of the leaf proteome of WT and ME plants after 15 (15D) and 30 (30D) days of drought. The proteomes of the ME1, ME3, and ME4 lines were combined (ME proteome). The 2,606 proteins identified in the 15D and 30D leaf samples (Dataset S2) were analysed for the identification of differentially expressed proteins (DEPs) using a *p*≤ 0.05 as the significance threshold parameter.

At 15D, we identified 109 proteins that increased (15DMEup cluster) in abundance in the ME lines compared to WT and 20 proteins that decreased (15DMEdown cluster) (Dataset S3). Functional enrichment analyses of these DEPs show that 20 proteins from the 15DMEup cluster are involved in photosynthesis, most of which are localized in the plastid (Fig. 5A, Dataset S3). These proteins are involved in light harvesting and electron transport (8 proteins; Fig. 5B) as well as in carbon fixation (12 proteins, of which 10 are involved in the Calvin cycle and two are phosphoenolpyruvate carboxylases; Fig. 5B). The 15DMEup cluster is also enriched for proteins involved in primary metabolism and translation (Fig. 5A). The 15DMEdown cluster is enriched for proteins involved in protein folding, ATP metabolism, and purine nucleotide metabolism (Fig. 5C).

**Fig. 5.**
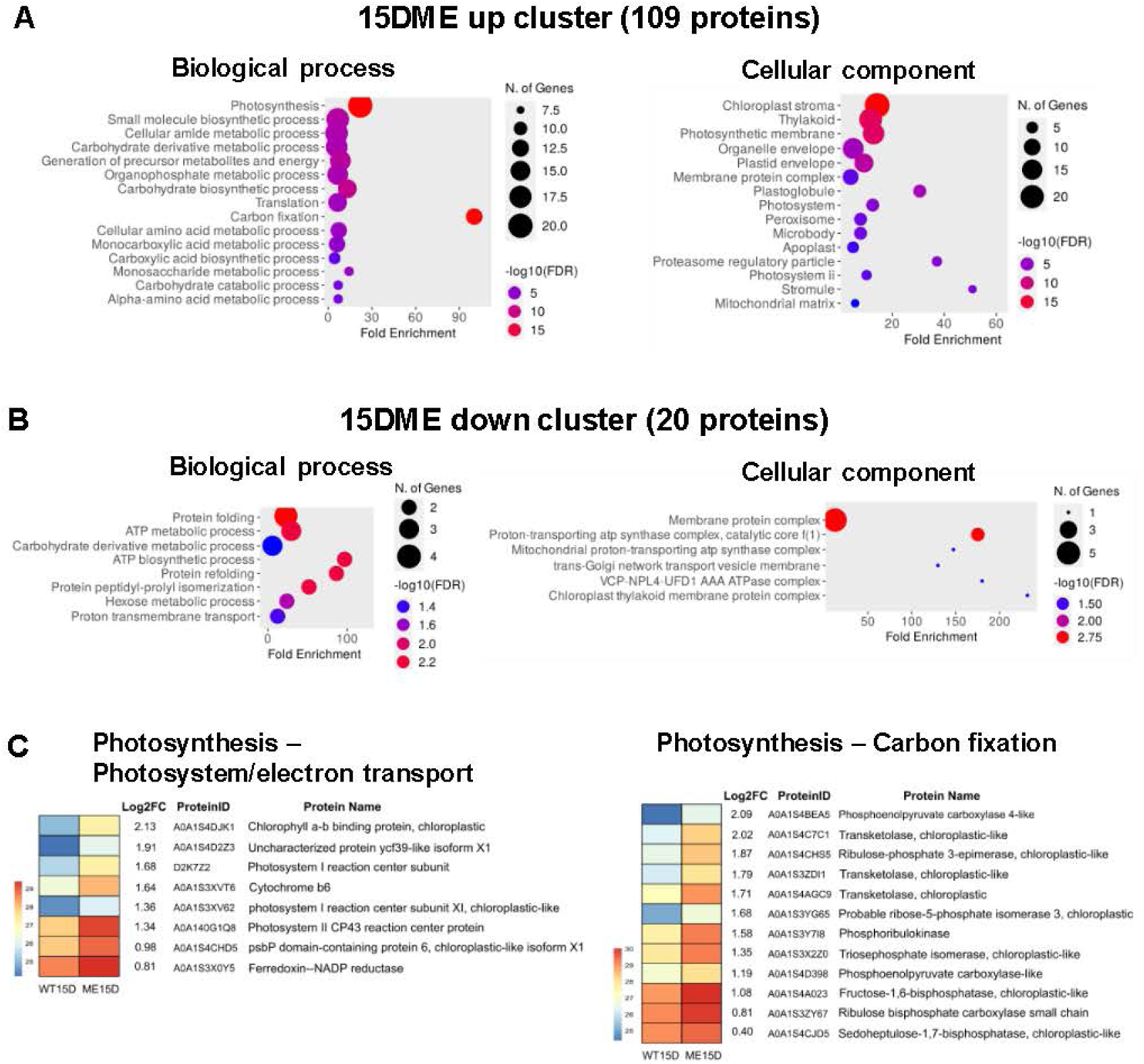
Differentially expressed proteins in leaves of ME and WT plants after 15 days of drought. Enriched biological processes and cellular components among the 109 more abundant proteins **(15DME up, A)**, and 20 less abundant proteins **(15DME down, B)** in ME lines compared to WT at 15D. The size of the circles indicates the number of proteins in each GO term, and the different colors indicate the –log10(FDR) values. **C.** Increased photosynthetic proteins in ME lines at 15D involved in photosynthetic electron transport and carbon fixation. The colored boxes indicate the log2 values of the protein intensities (a scale bar is included for each panel). Lists of the DEPs between ME lines and WT at 15D and the complete list of biological process, molecular function, and cellular component term enrichments among the DEPs can be found in Dataset S3.

At 30D, there are fewer DEPs between WT and ME lines than at 15D; 21 proteins increased in abundance (30DMEup cluster) and 22 decreased (30DMEdown cluster) in ME lines compared to WT (Dataset S3). The 30DMEup cluster was enriched in proteins located in plastids and related to carbohydrate metabolism and translation (Fig. S2); we found no enrichment of biological process terms in the 30DMEdown cluster (Dataset S3).

To identify differences in the proteomic response to drought between the genotypes, we searched for DEPs in the WT and ME between 15D and 30D. We found 119 DEPs in the WT, of which 95 were highly- and 24 less expressed between 15D and 30D (Dataset S4). We found 186 DEPs in the ME plants, of which 87 were highly- and 99 less expressed between 15D and 30D (Dataset S4). A Venn diagram of the DEPs shows that few DEPs overlap between the WT and ME plants; only 10 common proteins decrease in abundance and 17 increase between 15D and 30D, while 3 proteins show opposite responses in the WT (increase in abundance) and ME (decrease in abundance) plants (Fig. S3A, Dataset S4). Proteins related to photosynthesis were the most enriched in DEPs that decreased between 15D and 30D in both WT and ME lines (Fig. S3B, Dataset S4). However, more photosynthesis-related proteins were reduced in abundance in ME lines than in WT (24 out of 86 in ME lines, 3 out of 14 in WT, and 3 out of 10 common proteins were reduced in abundance in WT and ME), which is consistent with the fact that ME lines have a higher abundance of photosynthetic proteins at 15D than WT (Fig. 5B). We found some common GO term enrichments in WT and ME plants for proteins that increase in abundance between 15D and 30D; however, the abundance of only 17 common proteins were increased in both WT and ME lines, mainly related to detoxification and protein (un)folding processes (Fig. S3C, Dataset S4).

## Discussion

We found that tobacco lines expressing a maize plastidic NADP-ME under the KAT1 promoter show an increased seed yield under normal irrigation and after drought (Fig. 2), which additionally occurs much faster than in the WT (Table S1). Importantly, these lines maintain a high seed yield after recovery from 45 days of drought, a condition that the WT does not survive (Fig. 1 and 2).

One key aspect of the successful performance of the ME lines is that their low stomatal conductance (Fig. 4A) does not negatively affect their photosynthetic performance (Table S7). Moreover, after 30 days of drought, the drastic decrease in the stomatal conductance of the ME lines and the WT to 30% and 15% of the value at 90% FC, respectively (Fig. 4A), has different effects on their photosynthetic efficiency: while the ME lines use 50% of the energy absorbed by PSII for photochemical reactions, the WT uses only 25%. This improved photosynthetic performance of the ME lines correlates with an enrichment of proteins involved in the light-driven photosynthetic reactions and carbon assimilation compared to the WT under drought (Fig. 5A and 5B). The lower leaf temperature at 30D in the ME lines compared to the WT (Fig. 4E) may be due to their lower NPQ than the WT (Fig. 4C) and/or to an enhanced cooling effect due to morphological changes in their stems, as detailed below (Fig. 3). Overall, the WT responds to stomatal closure as expected (10, 28, 29), with reduced photochemical quenching and high leaf temperatures due to reduced cooling associated with reduced transpiration (Fig. 4A and 4C). In contrast, the ME lines break the trade-off between stomatal conductance, photosynthesis, and leaf temperature. The low stomatal conductance, which is usually an early response to drought, is a pre-drought state in the ME lines that establishes a new “homeostasis” at 90% FC. Thus, the ME lines are already “adapted” at 90% FC to maintain a high photosynthetic capacity despite their low stomatal conductance.

The question is how ME plants overcome the trade-off between stomatal conductance and photosynthesis. To answer this question, we must consider that tobacco, like other C_3_ species, has characteristics of C_4_ photosynthesis in cells surrounding the vascular tissues of stems and petioles, and that these cells are supplied with carbon for photosynthesis from the vascular system rather than from stomata (30). Therefore, it is likely that the introduction of a catalytically efficient NADP-ME (19, 31) into the cells close to the vascular system (Fig. 4D) improves the efficiency of this C_4_-like pathway by providing more CO_2_ from malate for sugar production despite stomatal closure. The increase in PEP-carboxylase, the primary carboxylating enzyme of C_4_ metabolism, in ME lines both at 90% FC (21) and after drought (Fig. 5B) supports this interpretation.

The transgenic NADP-ME activity is not restricted to vascular cells. In guard cells, this activity is probably related to the observed ABA-independent low stomatal conductance (Table S6). Stomata closure occurs due to a decrease in guard cells turgor induced by a decrease in K^+^ and anions such as Cl^−^ and malate (32, 33). However, malate not only plays a role as a counterion to K^+^ in guard cells, but also modulates the activity of several ion transporters, such as the plasma membrane S-type anion channel (34) and the vacuolar aluminum-activated malate transporter 9 (ALMT9) (18). Interestingly, ALMT9 is not only found in guard cells, but it is also highly expressed in the vasculature of shoots, leaves, and roots, where it regulates the expression of membrane transport proteins, such as the sucrose transporter (SUC) (35). Considering that SUC expression is increased in ME lines and induces higher leaf sugar export (21), we can hypothesize that the transgenic NADP-ME activity in cells close to the vasculature not only produces more CO_2_ for carbon fixation, but also induces an increase in SUC via a change in ALMT9 activity. Given that plant growth and yield ultimately depend on the amount of sucrose delivered to sink organs per unit of time (36, 37), the increase in carbon fixation and in sugar export in the ME lines is likely responsible for their increased seed yield (Fig. 2) and the production of seeds in a shorter time than in the WT (Table S1).

An unexpected phenotype of the ME lines is the morphology of their stems (Fig. 3A and 3B), which – as detailed below – may contribute to their rapid recovery from drought after rewatering (Fig. 1), their enhanced leaf cooling at 30D (Fig. 4E), and their increased height (21). Considering that adhesion forces between water and cell walls are higher in small than in large diameter vessels, leading to improved maintenance of the water column (38), the high proportion of narrow xylem vessels in ME lines (Fig. 3A and 3B) may result in more efficient water transport than the WT, reducing the likelihood of stem cavitation. It has also been shown that blocking sucrose loading and lack of sucrose in phloem sieve elements impairs xylem recovery from drought in tobacco (39). Thus, the efficient C_4_-like biochemistry and the increased sugar export of the ME lines may be critical for their rapid recovery from drought (Fig. 1). The increased growth rate (Table S1) and internodal length (21) of the ME lines may be related to their higher pith proportion in the stems, as plant species with the largest pith grow faster (40). The pith is a metabolically less active, low density tissue that is used for water storage in some succulent species (41). Apart from this, the exact physiological role of the pith in non-succulent plant species has received relatively little attention. We found only one study in tomatoes where the pith was disrupted in response to drought (stem pithiness) in an ABA-induced process (42).

Malate is an important carbon storage molecule with diverse signaling functions (43–49), as well as the products of the NADP-ME reaction, pyruvate and CO_2_ (50–52). Thus, it is not surprising that the constitutive plant-wide expression of NADP-ME in tobacco not only resulted in reduced stomatal conductance but also caused penalties in photosynthesis and growth (53). Our results show that a targeted expression strategy avoids such unwanted pleiotropic effects, demonstrating that in-depth knowledge of specific cellular metabolic processes can help develop precision metabolic engineering to improve crop performance and resilience in a changing climate.

## Materials and Methods

### Plant material and growth conditions

Experiments were conducted using *Nicotiana tabacum* (L. cv. Petit Havana SR1) wild-type (WT) and three independent transgenic homozygous lines (ME1; ME3; and ME4) expressing maize plastidic NADP-malic enzyme (AY315822) under the *Arabidopsis thaliana* Potassium channel 1 (KAT1) promoter (21). Plants were grown from seeds in a compost:sand:perlite mixture (2:1:1 by volume). Seedlings were transferred to a growth chamber kept at 30/18°C with 12/12 h day/night period and 200 μmol quanta m^−2^ s^−1^ of photosynthetic active radiation (PAR).

Flowering time was recorded as the time of growth of each plant until setting the first flower buds. Life cycle time was recorded as the time of growth until the plants produced mature seeds, after which the plants began the senescence process.

### Drought and recovery experiments

For the drought experiments, tobacco WT and transgenic M1, M3, and M4 lines were grown for 60 days in 3-L pots irrigated at 90 % field capacity (FC) (21). Fifteen 60-day-old plants of each genotype (WT, M1, M3, and M4) were exposed to a gradual increase in drought conditions by stopping the irrigation. In each experiment, 5 plants of each genotype were kept at 90 % FC irrigation (control plants) until completing their life cycle. Physiological and biochemical analyses were performed at different time points after drought (D samples), which were compared to control plants. Recovery from drought experiments was performed after 30 or 45 days without irrigation (30D and 45D plants, respectively) by adding water to the pots to 90% FC. During recovery, plants were analyzed at different time points until completing their life cycle. The samples from the drought-recovery experiment were named depending on the time of drought application and time of rewatering, for example, 30D3R or 30D7R for plants subjected to 30 days of drought followed by 3 or 7 days of recovery, respectively, and 30D+R for plants at the end of the life cycle. Pots containing WT and transgenic lines were randomly distributed in the same growth chamber. The drought and recovery experiments were repeated 4 different times (30D) and 3 different times (45D) from 2018 to 2021. The results obtained in the different experiments were comparable.

### Vegetative aerial biomass, seed yield and Harvest Index

Vegetative aerial biomass and fruit and seed production was analysed at the end of the life cycle of control plants (watered to 90% FC) and of plants recovered from 30D or 45D (30D+R and 45D+R, respectively) using at least three different plants per genotype and condition. For aerial vegetative biomass evaluation, leaves and stems were harvested and weighted after drying the plants in an oven at 60 °C for a week (dry weight, DW).

For reproductive biomass evaluation, the total number of capsules per plant were counted and collected at maturity. The seeds per capsule were collected and weighed separately. For seed number analysis, digital images of the seeds of four capsules of each genotype and condition were captured and processed using Image J software (54) to compute the number of seeds in the sample. 1000-seed weight was calculated by dividing seed weight by seed number and multiplying by 1000. The seed number per plant was calculated by dividing the total seed weight per plant by the average seed weight of each genotype. The harvest index (HI) was calculated as the ratio of seed yield (g) per plant to dry vegetative aerial plant biomass (leaves and stem without fruits, g) using at least three individual plants from each genotype.

### Seed composition and size measurement

Moisture content was determined weighing seeds from each genotype and condition before (fresh weight, FW) and after drying (dry weight, DW) at 80 °C to constant weight. For seed size measurement, mature seeds were photographed using a magnifying glass (Nikon). The diameter (mm) and area (mm^2^) of 100 seeds originated from three plants of each genotype and condition were measured with the Image J software (54). Starch was extracted from 30 mg of seed flour for 45 min at 95°C with KOH. After adjusting pH with acetic acid, the samples were incubated with amylase and amyloglucosidase for 16 h at room temperature. Samples were centrifuged at 15,000 g for 15 min and glucose was quantified in the supernatant by the glucose oxidase method using a commercial kit. Protein content of seed flour was measured by Kjeldahl method using the conversion factor of 6.25. Triglycerides (TAG) were extracted with ethanol at 50°C for 16 h (1:40 ethanol: seed powder). The enzymatic method (Triglyceride kit TG Color GPO/PAP AA, Wiener Lab) was used for TAG quantification.

### Extraction of stem RNA, in situ determination of ME activity and morphological studies

Stem samples at approximately 5 cm above the soil surface were collected from at least 3 plants of each genotype grown for 60 days at 90 % FC (control) and after 30 days of drought treatment (30D).

RNA was extracted from 100-150 mg of stem tissue from WT and ME lines using the protocol described in (55). For *ZmnpNADP-ME* transcript detection, RT-PCR studies were performed using the following primers: 5’CCATGGCTTCCTTCAATG3’and 5’CCGAACCAGGGAAATG’3, which amplify a 230-bp fragment of maize non photosynthetic NADP-ME (31). *In situ* NADP-ME activity assay was carried out in cross sections of WT and ME lines stems from 60-day-old tobacco plants as described in (56). For morphology studies, stem pieces of about 1 cm in length were fixed in FAA solution (50 % ethanol, 5 % glacial acetic acid, 30 % formaldehyde, 15 % water). Samples were dehydrated with ethanol and ethanol/xylene mixtures of increasing concentration and embedded in paraffin. Stem cross sections of 10 μm thickness were cut with a Minot microtome and stained with Safranin and Fast green. Sections were observed under light microscope (Zeiss MC 80 Axiolab), and the images were captured by a Nikon N11 camera and analysed using ImageJ software package (54). For control and 30D samples, the major cross-sectional diameter of the entire stem and pith; the major width of xylem tissue; and the area of the stem, pith, and xylem, were measured from at least 3 different sections of each sampled stem using ImageJ software package. The % of pith and xylem per stem area was calculated. Xylem vessels were classified depending on diameter length (0-20 µm, 20-40 µm, 40-60 µm, and 60-80 µm), and the number of vessels per size class were scored using at least 15 xylem vessels per image.

### Stomatal conductance and chlorophyll fluorescence measurements

Stomatal conductance (g_s_, mmol H_2_O m^−2^ s^−1^) of WT and ME lines was measured using a leaf porometer (SC-1 Porometer, Decagon Devices) between 4 to 6 h after the lights were turned on and using the fifth or sixth leaf from three to five plants of each genotype.

A MultispeQ v2.0 sensor (57) controlled by the PhotosynQ platform (Project ID 7925) was used to measure chlorophyll fluorescence quenching and other leaf traits (Tables S7 and Dataset S1). Measurements were performed using the “[Photosynthesis RIDES 2.0](https://www.photosynq.org/protocols/photosynthesis-rides-2-0)” protocol on leaf 5 of 60-day-old plants at 90% FC (before water stress), and on multiple leaves at 30D (leaf 6 to 10, numbered from bottom). The measurements were performed using at least three plants of each genotype between 4 to 6 h after illumination with a 200 μmol photons m^−2^ s^−1^ red actinic light. The minimal fluorescence in the light-adapted state (F_0_′) was determined by a short exposure (2 s) to weak far-red illumination (730 nm) without background actinic light illumination to oxidize the plastoquinone pool and Q_A_ while steady-state fluorescence (F_s_) was measured before the next flash. Saturating pulses of 6,000 μmol photons m^−2^ s^−1^ and 0.5 s duration were used to obtain maximal fluorescence signals (F_m_’). The resulting chlorophyll a fluorescence traces were used to estimate the following parameters: Quantum yield of photosystem (PSII) (Φ_PSII_ = (F_m_’ - F_s_)/F_m_’) (58), non-regulatory energy dissipation (Φ_NO_ = F_s_/F’_m_), and quantum efficiency of non-photochemical quenching (Φ_NPQ_ = 1 – Φ_NO_ – Φ_PSII_) (59) for light-adapted leaves; and (theoretical) non-photochemical quenching (NPQ_(T)_ = [4.88/(F’_m_/F’_0_)-1)]-1)) as defined in (60). Linear electron flow (LEF = Φ_PSII_ x PAR x 0.4); relative chlorophyll content (SPAD); leaf *versus* ambient temperature differential (δ Temp); and other leaf and environmental parameters (Table S7 and Dataset S1) were measured as described (57).

### Sample collection for the measurement of ABA levels

WT, ME1, and ME3 plants were grown from seed in soil. After 5 days of germination, the seedlings were grown in 5L pots in a growth chamber under 28/18 °C 12/12 h day/night photoperiod, 250 μmol quanta m^−2^ s^−1^ of PAR and irrigation at 90% FC. Stomatal conductance was measured for the 6th leaf of 35-day-old plants after 4-6 h of illumination using a LI-600 Porometer (Licor, Germany). Ten plants from each line were analysed. After 3 days of continuous stomatal conductance determination, the leaf tissue from the 6th leaf was collected after 4-6 h of illumination. One replicate of leaf tissue (100 mg) was collected from each plant for phytohormone determination and a total of 8 biological replicates were collected from each genotype. Samples were immediately frozen in liquid nitrogen and stored at −80°C. Watering was then stopped, and after seven days after drought (7D), the stomatal conductance of the 10th leaf was measured after 4-6 h of illumination. Samples were collected from the 10th leaf after 4-6 h of illumination for ABA measurement as indicated above.

### Tissue extraction for the measurement of ABA levels

Frozen leaves were ground with mortar and pestle. 50 mg plant material were used for extraction of ABA in a protocol adapted from (61). 500 µl of extraction solution (isopropanol/H_2_O/HCl (2:1:0.002; v/v/v), containing 0.0075 nmol of deuterated internal standard (D6-*cis*,*trans*-ABA, OlChemIm, Czech Republic) were added to each sample, followed by rigorous mixing and extraction of ABA in an ice-cold ultrasound bath for 30 min. In a second extraction step, 1 mL dichloromethane was added to each sample, followed by vortexing and incubation in an ice-cold ultrasound bath for 30 min. Phase separation was achieved by centrifugation at full speed (21.000 g, 4°C) for 5 minutes. The lower organic phase containing ABA was transferred to a fresh tube and dried under a stream of nitrogen. ABA was redissolved in 50 µl MeOH, 50 µl H_2_O was added and the samples kept at 4°C overnight and centrifuged for 5 minutes at full speed (21.000 g, 4 °C) to remove residual undissolved precipitated compounds. The resulting supernatant was used for LC-MS analysis of ABA (See SI Appendix for details.

### Sample collection for proteome analysis and quantification by LC-MS

Total protein was extracted from the seventh leaf of the WT and ME transgenic lines after 15 (15DWT and 15DME samples) and 30 days without watering (30DWT, and 30DME samples) as previously described (Fig. 1). Three biological replicates (a, b, and c) for each WT sample (15DWT and 30DWT), and two biological replicates (a and b) for each ME transgenic line sample (15DME1, −3, −4, and 30DME1, −3, and −4) were analysed, resulting in a total of 18 samples. Total protein from each sample was extracted, quantified, treated, and trypsin-digested, and desalted as in (21). The peptides obtained were resolved in a nano-HPLC (EASY-nLC 1000, ThermoScientific, Germany), ionized by electrospray, and analysed by a mass spectrometer (MS) with Orbitrap technology (Q-Exactive with High Collision Dissociation cell and Orbitrap analyser, ThermoScientific, Germany) at the Proteomics Core Facility CEQUIBIEM (CONICET, Buenos Aires University). Protein identification was performed with Proteome Discoverer 2.1 software (ThermoScientific, Germany) using *N. tabacum* Uniprot proteome as reference (https://www.uniprot.org/taxonomy/4097). Protein hits with at least two peptides detected were filtered for high confidence peptide matches, resulting in 2,606 identified proteins. Dataset S2 lists the accession number in UniprotKB database (http://www.uniprot.org/uniprot/); coverage and peptides used for identification; area for each biological repetition; as well as other parameters (such as Molecular Weight and calculated Isoelectric Point) of the 2,606 identified proteins.

### Differential proteome analysis

Proteomic data (Dataset S2) was analysed with Perseus software (Max Planck Institute of Biochemistry, http://www.perseus-framework.org) (62). Differential proteome analysis was performed by comparing 15DWT versus 15DME, 30DWT versus 30DME, 15DWT versus 30DWT, and 15DME versus 30DME values. In all cases, proteomic results from the three ME lines (1, 3, and 4) were pooled together and considered as if they were derived from a unique ME genotype, resulting in six biological replicates for 15DME and 30DME samples. Normalised label-free quantification (LFQ) protein intensities were log2 transformed and filtered for at least two valid values among the three biological replicates of 15DWT and 30DWT, and at least four replicate valid values of 15DME and 30DME. Statistical analysis (Student’s t-test) was performed using the Perseus 1.6.5.0 software. A p-value ≤ 0.05 was used as significance threshold parameter in all cases to identify differentially expressed proteins (DEP).

DEP increasing (up) or decreasing (down) in ME lines with respect to WT after 15 and 30 days without watering are listed in Dataset S3 as follows: 15DMEup, 15DMEdown, 30DMEup, and 30DMEdown. DEP increasing (up) and decreasing (down) by drought treatment (from 15D to 30D) in WT and ME lines are listed in Dataset S4.

Protein lists were submitted to a gene ontology (GO) term enrichment analysis using ShinyGo 0.80 (http://bioinformatics.sdstate.edu/go80/) (63). Biological process (BP), molecular function (MF), and cellular component (CC) term enrichment was requested for each protein list using *Nicotiana tabacum* data as available from ShinyGo tool (STRINGv11.5). Venn diagram for DEPs of WT and ME lines from 15D to 30D was generated using Venny 2.1 tool (https://bioinfogp.cnb.csic.es/tools/venny/).

### Statistical Analyses

Except for proteomic analysis where Perseus software was used, statistical computing was performed with R software. Statistical differences of normal data were determined using one-way ANOVA and Tukey’s multiple comparison test (*p*<0.05). Shapiro-Wilk tests were used to verify the normal distribution of the data; a *p*-value below 0.05 was taken as indication that the data was non-normal. In the case of non-normal data, the Kruskal– Wallis test and the pairwise Wilcoxon Rank Sum test were used (*p*< 0.05).

## Author Contributions

MFD, GLM, MVL, and VGM conceived research, and supervised data analysis and interpretation. PO, MFG, GLM, MSS, CS, YF, and KV performed research and analysed data. EM and GME performed proteomic and photosynthetic data analysis and interpretation. SM and VW contributed with analytical tools. MFD, VGM, and GME wrote the manuscript. All authors contributed to the generation of the figures and the revision of the manuscript. All authors accepted the final version of the manuscript.

## Acknowledgments

This work was funded by grants of the Agencia Nacional de Promoción Científica y Tecnológica (PICT RAICES 2018-0813 and PICT 2021 CAT I 0006, Argentina); and the Deutsche Forschungsgemeinschaft (DFG, German Research Foundation) under Germanýs Excellence Strategy – EXC-2048/1 - project ID 390686111. MFD, MFG, GLM, EM, and MVL are members of the Researcher Career of CONICET. Part of the results presented are described in the patent AR072885B1 “Método para inducir el crecimiento y/o floración temprana de las plantas” (www.inpi.gob.ar).

