## Supplemental Information for "Drought resistance and improved yield result from modified malate metabolism in guard and vascular companion cells"

**This PDF file includes:**

Supporting text
Figures S1 to S3
Tables S1 to S7
Legends for Datasets S1 to S4

**Other supporting materials for this manuscript include the following:**

Datasets S1 to S4

### Supporting Information Text

#### *LC-MS analysis of ABA*

10 µl sample volumes of ABA extracts were used for injection. ABA isomers were separated on a Dionex HPG 3200 HPLC system (Thermo Scientific, Germany) equipped with a C18 XSelect® HSS T3 Column, 100Å, 2.5 µm, 3 mm X 150 mm (Waters, Ireland) with a binary gradient system. Mobile phase A consisted of water + 0.1% formic acid (FA) and mobile phase B consisted of methanol + 0.1% FA. The mobile phase gradient was as follows: Starting conditions were 2% mobile phase B, increased to 99% B from 1-18 min, the plateau was held for 4 min and the system was returned to starting conditions within 1 min. The system was kept at 2% B for another 7 min for equilibration prior to the next injection. The flow rate was 0.35 ml/min. ABA isomers were analyzed by Q-TOF MS on a maXis 4G instrument (Bruker Daltonics, Germany) equipped with an ESI source. The instrument was operated in negative-ion mode and the operating conditions were as follows: dry gas (nitrogen): 8.0 l/min, dry heater: 200 °C, nebulizer pressure: 1.0 bar, capillary voltage: 4500 V. Data were acquired in stepping mode with collison Rf voltage ranging from 300 to 500 Vpp. (+)-*trans*, *trans*-ABA and (+)-*cis*, *trans*-ABA isomers eluted at 14.2 min and 14.8 minutes and were quantified in relation to a deuterated internal standard (D6-*cis*, *trans*-ABA) using calibration curves obtained with commercial reference standards (+)-*trans*, *trans*-ABA (OlChemIm, Czech Republic, cat.no. 0132751); (±)-*cis*, *trans*-ABA, (OlChemIm, Czech Republic, cat.no. 0132701). D6-(+)-*cis*, *trans*-abscisic acid (D6-ABA (OlChemIm, Czech Republic, cat.no. 0342721) was used as an internal standard.

### SI Figures

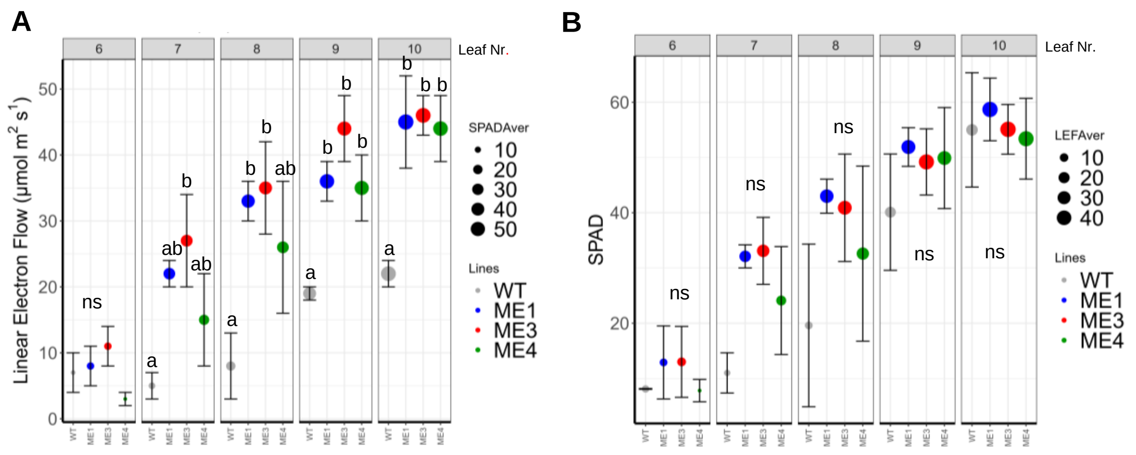


**Fig. S1. Chlorophyll fluorescence measurements of WT and ME lines after 30 days of drought.** Measurements were performed on individual leaves 6 to 10, numbered from bottom to top, from three different pants of each genotype using the MultispeQ v2.0 sensor (PhotosynQ platform Project ID 7925). The leaf number (6 to 10) is indicated at the top of each plot. **A.** Linear electron flow. **B.** Relative chlorophyll content (SPAD). Different letters indicate significant differences (*p* value <0.05) according to Tukey´s test of ANOVA. ns: not significant.

### 30D ME up cluster (21 proteins)

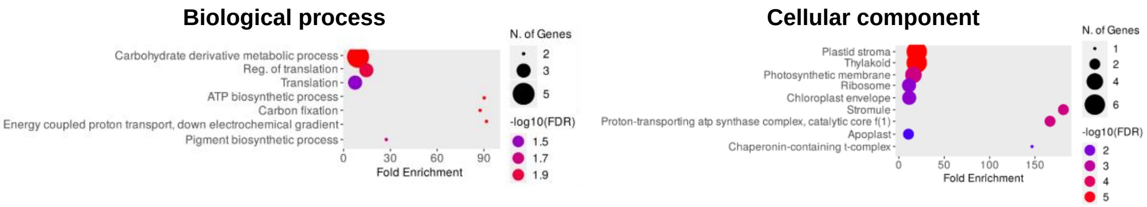


**Fig. S2.** Representative enriched biological processes and cellular components among the 21 more abundant proteins (30DMEup) in ME lines compared to WT at 30D. The size of the circles indicates the number of proteins in each GO term, and the different colors the –log10(FDR) values. A list of the DEPs between ME lines and WT at 30D and the complete list of biological process, molecular function, and cellular component term enrichments among the DEPs can be found in Dataset S3.

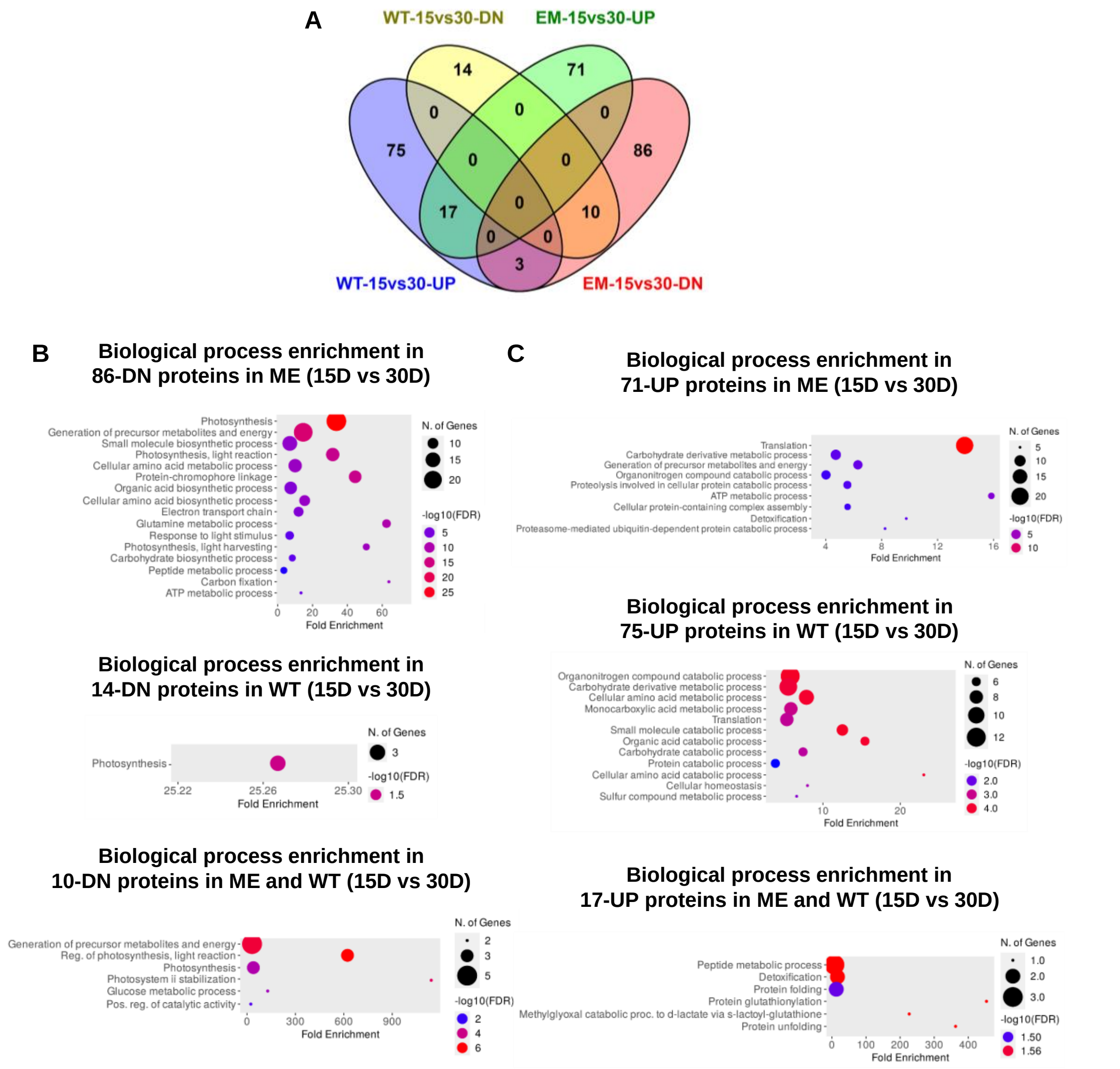


**Fig. S3. Differentially expressed proteins in leaves of ME and WT plants after 15 and 30 days of drought. A.** Venn diagram showing the number of DEPs (DN, decreased; UP, increased) after comparing 15D with 30D in the WT and ME lines independently. **B** and **C**. Enriched biological processes among the proteins decreased (DN, B) or increased (UP, C) only in WT or in ME lines, or in both genotypes. The size of the circles indicates the number of proteins in each GO term, and the different colors indicate the –log10(FDR) values. A list of the DEPs increasing or decreasing in WT or in ME lines, or in both WT and ME lines, and the biological process, molecular function, and cellular component GO term enrichment analysis of each cluster can be found in Dataset S4.

### SI Tables

**Table S1. Flowering time and life cycle duration of plants at 90 % FC and after recovery from 30 days of drought.** Tobacco WT and ME lines were grown at 90% FC. After 60 days of growth, one group of plants was exposed to 30 days of drought followed by rewatering at 90% FC (30D+R), while the other group was watered normally (control). Flowering time (time of growth until setting of the first flower bud) and the life cycle (time of growth until production of mature seeds) were calculated for at least 10 plants of each genotype. Different letters indicate significant differences ($p<0.05$) after Tukey´s test of ANOVA (life cycle) or Kruskal-Wallis test (flowering time).

| | WT | ME1 | ME3 | ME4 |
|---|---|---|---|---|
| | **90% FC (control)** | | | |
| **Flowering time (days)** | 98.9 ± 4.1 b | 71.5 ± 0.9 a | 71.4 ± 1.4 a | 71.3 ± 2.0 a |
| **Life cycle (days)** | 147.3 ± 3.3 b | 116.9 ± 1.6 a | 118.1 ± 1.5 a | 117.3 ± 1.3 a |
| | **Recovery after 30 days of drought (30D+R)** | | | |
| **Flowering time (days)** | 122.5 ± 4.5 b | 99.0 ± 1.0 a | 101.5 ± 1.5 a | 98.5 ± 0.5 a |
| **Life cycle (days)** | 150.0 ± 5.0 b | 127.5 ± 0.5 a | 131.0 ± 2.0 a | 129.5 ± 0.5 a |

**Table S2. Vegetative aerial biomass at 90% FC and after recovery from 30 and 45 days of drought.** Tobacco WT and ME lines were grown at 90% FC (control) or exposed to 30 or 45 days of drought followed by recovery by rewatering at 90% FC. At the end of the life cycle (30D+R and 45D+R), the plants were harvested, and the dry weight (DW) of the leaves and stems of each plant was measured. The average DW ± SD of 3 to 5 plants of each genotype and the sum of these averages (total vegetative aerial DW) are given. Different letters indicate significant differences (p value <0.05) after the Tukey´s test of ANOVA or the Kruskal-Wallis test (leaves DW at 90% FC; stem DW at 30D+R; and total vegetative weight at 30D+R). N.R. (No Recovery): WT plants do not recover after 45D (Fig. 1B).

| | WT | ME1 | ME3 | ME4 |
|---|---|---|---|---|
| | **90% FC (control)** | | | |
| **Leaves DW (g)** | 6.9 ± 2.6 a | 4.3 ± 0.1 a | 3.7 ± 0.4 a | 4.3 ± 0.9 a |
| **Stem DW (g)** | 5.6 ±0.8 b | 4.2 ± 0.7 ab | 3.5 ± 0.6 a | 3.7 ± 0.9 a |
| **Total vegetative aerial DW (g)** | 12.5±3.5 b | 8.5±0.8 a | 7.2±0.9 a | 8.0±1.9 a |
| | **Recovery after 30 days of drought (30D+R)** | | | |
| **Leaves DW (g)** | 5.1 ± 0.1 b | 3.2 ± 0.2 a | 3.2 ± 0.2 a | 3.3 ± 0.7 a |
| **Stem DW (g)** | 4.8 ± 1.2 b | 2.8 ± 0.3 ab | 2.3 ± 0.2 a | 2.7 ± 0.2 ab |
| **Total vegetative aerial DW (g)** | 9.4 ± 1.9 b | 6.0 ± 0.1 ab | 5.5 ± 0.1 a | 6.0 ± 0.9 a |
| | **Recovery after 45 days of drought (45D+R)** | | | |
| **Leaves DW (g)** | N.R. | 3.4 ± 0.8 a | 4.0 ± 0.5 a | 3.0 ± 0.3 a |
| **Stem DW (g)** | N.R. | 2.4 ± 0.4 a | 2.5 ± 0.5 a | 2.1 ± 0.6 a |
| **Total vegetative aerial DW (g)** | N.R. | 5.8 ± 1.2 a | 6.5 ± 0.9 a | 5.2 ± 0.9 a |

**Table S3. Reproductive biomass at 90 % FC and after recovery from 30 and 45 days of drought.** Tobacco WT and ME lines were grown at 90 % FC (control) or exposed to 30 or 45 days of drought followed by rewatering at 90 % FC (30D+R or 45D+R). Total reproductive biomass was measured in 2 independent biological repetitions with similar results, using at least 3 plants of each genotype. Results are from the experiment in 2021. Capsules were collected at maturity and the seeds per capsule were weighed separately. For seed number analysis, digital images of the seeds from four capsules were processed using Image J software. The 1000 seed weight was calculated by dividing the seed weight by the seed number and multiplying by 1000. Seed number per plant was calculated by dividing the total seed weight per plant by the average seed weight of each genotype. Harvest index was calculated as the ratio of the seed yield to the vegetative plant biomass (Table S2). The mean ± SD value obtained from at least 3 plants of each genotype is given. Different letters indicate significant differences (p value <0.05) after the Tukey´s test of ANOVA or the Kruskal-Wallis test (seed number per fruit at 30D+R; seed weight per fruit at 45D; and Harvest Index at 90 % FC and 30D+R). NR, no recovery of WT plants after 45D+R treatment (Fig. 1B)

| | WT | ME1 | ME3 | ME4 |
|---|---|---|---|---|
| | **90% FC (control)** | | | |
| **Fruit number per plant** | 28.3 ± 6.7 a | 42.7 ± 6.0 a | 34.0 ± 7.6 a | 39.3 ± 8.3 a |
| **Seed number per fruit** | 1246 ± 431 a | 2556 ± 608 b | 2394 ± 636 b | 2593 ± 238 b |
| **Seed number per plant** | 25801 ± 6423 a | 75338 ± 22322 b | 52732 ± 10204 ab | 55603 ± 20922 ab |
| **Seed weight per fruit (g)** | 0.10 ± 0.03 a | 0.13 ± 0.04 b | 0.13 ± 0.04 b | 0.14 ± 0.04 b |
| **Seed weight per plant (g)** | 1.96 ± 0.35 a | 3.45 ± 0.46 b | 2.73 ± 0.37 ab | 3.31 ± 0.48 b |
| **Harvest index** | 0.15 ± 0.08 a | 0.41 ± 0.04 b | 0.39 ± 0.07 b | 0.42 ± 0.05 b |
| **1000 seeds weight (g)** | 0.08 ± 0.01 b | 0.05 ± 0.02 a | 0.05 ± 0.01 a | 0.05 ± 0.01 a |
| | **Recovery after 30 days of drought (30D+R)** | | | |
| **Fruit number per plant** | 26.7 ±2.1 a | 31.5 ± 0.7 a | 30.0 ± 6.1 a | 31.7 ± 5.7 a |
| **Seed number per fruit** | 1175 ±267 a | 2324 ± 395 c | 2281 ± 298 c | 1835 ± 553 b |
| **Seed number per plant** | 24612 ± 4072 a | 48540 ± 603 c | 46266 ± 3098 c | 35633 ± 1633 b |
| **Seed weight per fruit (g)** | 0.09 ± 0.02 a | 0.13 ± 0.02 b | 0.12 ± 0.03 b | 0.12 ± 0.03 b |
| **Seed weight per plant (g)** | 1.88 ± 0.11 a | 2.74 ± 0.02 c | 2.36 ± 0.23 b | 2.3 ± 0.16 b |
| **Harvest index** | 0.20 ± 0.04 a | 0.46 ± 0.01 c | 0.43 ± 0.03 bc | 0.39 ± 0.03 b |
| **1000 seeds weight (g)** | 0.08 ± 001 c | 0.06 ± 0.01 ab | 0.05 ± 0.01 a | 0.07 ± 0.01 b |
| | **Recovery after 45 days of drought (45D+R)** | | | |
| **Fruit number per plant** | NR | 29.5 ± 7.8 a | 23.3 ± 4.2 a | 33.0 ± 7.1 a |
| **Seed number per fruit** | NR | 1471 ± 428 a | 2213 ± 708 b | 2324 ± 252 b |
| **Seed number per plant** | NR | 43896 ± 10605 a | 41526 ± 10999 a | 42584 ± 3899 a |
| **Seed weight per fruit (g)** | NR | 0.09 ± 0.03 a | 0.13 ± 0.03 b | 0.14 ± 0.01 b |
| **Seed weight per plant (g)** | NR | 2.7 ± 0.5 a | 2.3 ± 0.3 a | 2.6 ± 0.4 a |
| **Harvest index** | NR | 0.48 ± 0.18 a | 0.37 ± 0.1 a | 0.51 ± 0.01 a |
| **1000 seeds weight (g)** | NR | 0.06 ± 0.01 a | 0.06 ± 0.01 a | 0.06 ± 0.0046 a |

**Table S4. Seed composition and size at 90% FC and after recovery from 30 and 45 days of drought.** Tobacco WT and ME lines were grown at 90% FC (control) or exposed to 30 or 45 days of drought followed by rewatering at 90 % FC (30D+R or 45D+R). Seeds from at least 3 plants of each genotype and condition were collected and analyzed for size and composition. Mean ± SD are given. Different letters indicate significant differences (p value <0.05) after the Tukey´s test of ANOVA or the Kruskal-Wallis test (starch at 90% FC). TAG: Triacylglycerol. NR: no recovery of the WT plants after 45D (Fig. 1B).

| | **WT** | **ME1** | **ME3** | **ME4** |
|---|---|---|---|---|
| | **90% FC (control)** | | | |
| **Starch (mg per 100 g DW)** | 57.7 ± 16.0 a | 59.9 ± 11.2 a | 49.7 ± 3.7 a | 49.8 ± 4.5 a |
| **Protein (g per100 g DW)** | 20.8 ± 0.7 a | 19.6 ± 0.19 a | 19.3 ± 0.4 a | 19.7 ± 0.39 a |
| **TAG (g per 100 g DW)** | 14.1 ± 4.1 a | 16.9 ± 2.9 a | 18.3 ± 7.9 a | 20.4 ± 3.9 a |
| **Water content (%)** | 5.77 ± 1.51 a | 5.43 ± 0.33 a | 5.71 ± 0.07 a | 6.09 ± 1.13 a |
| **Seed diameter (mm)** | 0.77 ± 0.01 a | 0.75 ± 0.02 a | 0.72 ± 0.01 a | 0.74 ± 0.02 a |
| **Seed area ($mm^2$)** | 0.34 ± 0.02 b | 0.31 ± 0.02 ab | 0.29 ± 0.02 a | 0.31 ± 0.02 ab |
| | **Recovery after 30 days of drought (30D+R)** | | | |
| **Starch (mg per 100 g DW)** | 70.5 ± 12.9 a | 58.0 ± 6.8 a | 72.8 ± 14.3 a | 58.1 ± 9.9 a |
| **Protein (g per100 g DW)** | 21.9 ± 0.6 a | 19.2 ± 0.86 a | 19.9 ± 0.3 a | 20.7 ± 0.53 a |
| **TAG (g per 100 g DW)** | 11.0 ± 1.7 a | 14.4 ± 0.3 a | 13.2 ± 3.7 a | 15.4 ± 2.9 a |
| **Water content (%)** | 5.21 ± 0.04 a | 5.12 ± 0.46 a | 6.03 ± 0.44 a | 5.15 ± 0.11 a |
| **Seed diameter (mm)** | 0.77 ±0.003 c | 0.74 ± 0.01 b | 0.72 ± 0.004a | 0.74 ± 0.01 b |
| **Seed area ($mm^2$)** | 0.34 ± 0.004 c | 0.30 ± 0.01b | 0.29 ± 0.003a | 0.30 ± 0.003b |
| | **Recovery after 45 days of drought (45D+R)** | | | |
| **Starch (mg per 100 g DW)** | NR | 63.7 ± 12.9 a | 63.5 ± 4.4 a | 52.5 ± 10.5 a |
| **Protein (g per100 g DW)** | NR | 20.4 ± 0.11 a | 21.2 ± 0.4 a | 20.2 ± 0 a |
| **TAG (g per 100 g DW)** | NR | 11.8 ± 7.9 a | 15.2 ± 6.1 a | 22.3 ± 1.0 a |
| **Water content (%)** | NR | 6.57 ± 0.76 a | 5.34 ± 0.37 a | 5.74 ± 0.64 a |
| **Seed diameter (mm)** | NR | 0.76 ± 0.03 a | 0.74 ± 0.01a | 0.75 ± 0.02 a |
| **Seed area ($mm^2$)** | NR | 0.32 ± 0.01 a | 0.30 ± 0.01 a | 0.32 ± 0.02 a |

**Table S5.** Analysis of stems at 90% FC and after 30 days of drought. Stem samples were taken at approximately 5 cm above the soil surface from at least 3 plants of each genotype grown for 60 days at 90% FC and after 30D. Light microscope cross-sections of the stems were analyzed using the ImageJ software package. The main cross-sectional diameter of the pith; the main width of the cortex and xylem tissue; and the area of the pith, xylem, and cortex were measured from at least 3 different sections of each sampled stem. The percentage of stem area occupied by the pith, xylem, and cortex is given in brackets. Different letters indicate significant differences (p value < 0.05) after the Tukey´s test of ANOVA or the Kruskal-Wallis test (cortex diameter at 90% FC and 30D; main xylem width, and stem, xylem, and cortex area at 30D).

| | WT | ME1 | ME3 | ME4 |
|---|---|---|---|---|
| | **60-day-old plants – 90% FC** | | | |
| **Main pith diameter (mm)** | 2.0 ± 0.1 a | 2.6 ± 0.1 b | 2.4 ± 0.1 b | 2.7 ± 0.3 b |
| **Pith area ($mm^2$)** | 2.3 ± 0.2 a (13%) | 2.8 ± 0.3 a (20%) | 4.4 ± 0.2 b (31%) | 3.8 ± 0.7 b (22%) |
| **Main xylem width (mm)** | 0.88 ± 0.13 c | 0.48 ± 0.05 b | 0.29 ± 0.05 a | 0.53 ± 0.09 b |
| **Xylem area ($mm^2$)** | 2.8 ± 1.0 a (15%) | 1.2 ± 0.2 b (9%) | 1.1 ± 0.1 b (8%) | 1.7 ± 0.2 b (10%) |
| **Major cortex diameter (mm)** | 2.3 ± 0.17 b | 2.6 ± 0.2 b | 1.6 ± 0.14 a | 2.4 ± 0.4 b |
| **Cortex area ($mm^2$)** | 12.8 ±1.3 c (72%) | 9.8 ± 0.92 ab (71%) | 8.8 ± 0.72 a (61%) | 11.6 ± 2.4 bc (68%) |
| | **30 days of drought (30D)** | | | |
| **Main pith diameter (mm)** | 2.2 ± 0.4 a | 4.3 ± 0.2 b | 4.0 ± 0.5 c | 3.3 ± 0.2 c |
| **Pith area ($mm^2$)** | 3.2 ± 0.9 a (6%) | 12.3 ± 2.2 b (24%) | 10.7 ± 3.3 c (22%) | 8.1 ± 0.7 c (14%) |
| **Main xylem width (mm)** | 3.4 ± 0.5 b | 1.9 ± 0.2 a | 2.4 ± 0.5 a | 2.9 ± 0.1 b |
| **Xylem area ($mm^2$)** | 18.5 ± 2.1 b (35%) | 14.3 ± 1.0 a (30%) | 15.7 ± 1.9 a (32%) | 18.0 ± 1.1 b (32%) |
| **Major cortex diameter (mm)** | 2.9 ± 0.2 b | 2.3 ± 0.2 a | 2.3 ±0.2 a | 2.9 ± 0.2 b |
| **Cortex area ($mm^2$)** | 30.6 ± 2.8 b (59%) | 21.9 ± 1.4 a (46%) | 22.3 ± 2.1 a (46%) | 30.5 ± 2.8 b (54%) |

**Table S6.** ABA levels in leaves of WT and ME plants. The levels of (+)-*cis, trans*-ABA and (+)-*trans, trans*-ABA and conductance were determined in leaf extracts of 38-day-old WT and ME plants grown at 90% FC and after 7 days of water withdrawal (7D). Samples were collected after 4-6 h of light. Mean ± SD of 8 independent samples of each genotype is given. Values significantly different ($p < 0.05$ as determined by a Student's *t* test) from the WT are shown in bold.

| | | **(+)-*cis, trans*-ABA** Average ± SD (ng $g^{-1}$FW) | p | **(+)-*trans, trans*-ABA** Average ± SD (ng $g^{-1}$FW) | p | **gs** Average ± SD (mol $m^{-2}$ $s^{-1}$) | p |
|---|---|---|---|---|---|---|---|
| **90% FC** | **WT** | 6.22 ± 3.17 | | 0.47 ± 0.14 | | 0.26 ± 0.04 | |
| | **ME1** | 5.33 ± 1.28 | 0.250 | 0.59 ± 0.22 | 0.111 | **0.16 ± 0.03** | **0.00002** |
| | **ME3** | 6.18 ± 1.14 | 0.466 | **0.69 ± 0.25** | **0.013** | **0.18 ± 0.02** | **0.00044** |
| **7D** | **WT** | 186.64 ± 126.77 | | 1.87 ± 0.63 | | 0.016 ± 0.036 | |
| | **ME1** | **65.19 ± 14.57** | **0.012** | 1.41 ± 0.22 | 0.123 | 0.029 ± 0.014 | 0.490 |
| | **ME3** | **99.96 ± 28.75** | **0.049** | 2.12 ± 0.55 | 0.114 | 0.027 ± 0.051 | 0.750 |

**Table S7**. Chlorophyll fluorescence parameters of 60-day-old WT and ME lines at 90% FC. Measurements were performed on leaf number 5 using the MultispeQ v2.0 sensor controlled by the PhotosynQ platform (Project ID 7925). Three 60-day-old plants at 90% FC from each genotype were used. Quantum yield of photosystem PSII (ΦPSII), non-regulatory energy dissipation (ΦNO), non-photochemical quenching (ΦNPQ), linear electron flow (LEF), relative chlorophyll content (SPAD), and the difference between leaf and ambient temperature (δ Temp) are shown. Different letters indicate significant differences ($p < 0.05$) according to Tukey´s test of ANOVA.

| | **WT** | **ME1** | **ME3** | **ME4** |
|---|---|---|---|---|
| | **60-day-old plants – 90% FC** | | | |
| **$\Phi_{PSII}$** | 0.61 ± 0.01 a | 0.59 ± 0.02 a | 0.59 ± 0.02 a | 0.61 ± 0.01 a |
| **$\Phi_{NPQ}$** | 0.17 ± 0.01 a | 0.19 ± 0.02 ab | 0.20 ± 0.02 b | 0.18 ± 0.01 ab |
| **$\Phi_{NO}$** | 0.21 ± 0.01 a | 0.22 ± 0.01 a | 0.21 ± 0.01 a | 0.21 ± 0.01 a |
| **LEF** | 53.5 ± 7.1 a | 55.6 ± 4.9 a | 54.9 ± 6.1 a | 54.5 ± 5.7 a |
| **SPAD** | 30.9 ± 2.9 a | 34.0 ± 2.3 a | 34.2 ± 1.2 a | 34.6 ± 2.0 a |
| **δ Temp** | -2.48 ± 3.53 a | -2.96 ± 1.59 a | -3.03 ± 1.91 a | -3.45 ± 2.22 a |

**SI Datasets**

**Dataset S1 (separate file).** Chlorophyll a fluorescence quenching and other leaf traits were measured using a MultispeQ v2.0 sensor controlled by the PhotosynQ platform (Project ID 7925). Leaves 6 to 10 (counting from bottom) from at least three different plants of each genotype were measured using the "Photosynthesis RIDES 2.0" protocol during the morning hours (between 4 to 6 h after illumination). The Table shows the results obtained for each individual plant measured: leaf versus ambient temperature differential ($\delta$ Temp); linear electron flow (LEF); theoretical non-photochemical quenching (NPQ(T)); quantum yield of PSII ($\Phi$PSII); non-regulatory energy dissipation ($\Phi$NO); quantum efficiency of non-photochemical quenching ($\Phi$NPQ); and relative chlorophyll content (SPAD).

**Dataset S2 (separate file). Proteomic data of *KAT1::ZmnpNADP-ME* lines after 15 and 30 days without watering.** Complete Database of identified proteins of the WT and ME transgenic lines M1, M3, and M4 after 15 (15DWT and 15DME1, -3 and -4 samples) and 30 days (30DWT and 30DME1, -3 and -4 samples) without watering. Triplicates from 15DWT and 30DWT, and duplicates from 15DME1, -3, -4, and 30DME1, -3, and -4 are denoted with a, b, or c after sample number (e.g. 15DWT a). The UniProtKB database (http://www.uniprot.org/uniprot/) accession numbers of each identified protein using the Proteome Discoverer software are indicated. For each identified protein, the coverage; number of peptides (#peptides); Peptide Spectrum Matches (#PSM); number of unique peptides (# Unique Peptides); number of AA composing (#AA); Molecular Weight (MW, kDa); calculated isoelectric point (calc. pI.); and Label-free quantification values as Areas of each biological replicate, are indicated.

**Dataset S3 (separate file). DEPs between WT and ME lines after 15 or 30 days of drought.** Differential proteome analysis was performed by comparing 15DWT versus 15DME, and 30DWT versus 30DME. DEPs are clustered as follows: 15DMEup, 15DMEdown, 30DMEup, and 30DMEdown, for proteins increasing (up) or decreasing (down) in ME lines with respect to WT after 15 or 30 days without watering, respectively. Biological process, molecular function, and cellular component GO term enrichment analysis of each DEP cluster is indicated.

**Dataset S4 (separate file). DEPs increasing or decreasing from 15D to 30D in WT and ME lines.** Differential proteome analysis was performed by comparing 15DWT versus 30DWT, and 15DME versus 30DME. According to Venn clustering, proteins increasing (UP) or decreasing (DN) only in WT or in ME lines, or in both WT and ME lines, are indicated. For each cluster, the biological process, molecular function, and cellular component GO term enrichment analysis is indicated.
